# Activation of cGAS/STING pathway upon paramyxovirus infection

**DOI:** 10.1101/2020.12.26.424443

**Authors:** Mathieu Iampietro, Claire Dumont, Cyrille Mathieu, Julia Spanier, Jonathan Robert, Aude Charpenay, Sébastien Dupichaud, Kévin P. Dhondt, Noémie Aurine, Rodolphe Pelissier, Marion Ferren, Stéphane Mély, Denis Gerlier, Ulrich Kalinke, Branka Horvat

## Abstract

During inflammatory diseases, cancer and infection, the cGAS/STING pathway is known to recognize foreign or self-DNA in the cytosol and activate an innate immune response. Here, we report that negative-strand RNA paramyxoviruses, Nipah virus (NiV) and Measles virus (MeV), can also trigger the cGAS/STING axis. While mice deficient for MyD88, TRIF and MAVS still moderately control NiV infection when compared to WT mice, additional STING deficiency resulted in 100% lethality, suggesting synergistic roles of these pathways in host protection. Moreover, deletion of cGAS or STING resulted in decreased type-I interferon production with enhanced paramyxoviral infection in both human and murine cells. Finally, the phosphorylation and ubiquitination of STING, observed during viral infections, confirmed the activation of cGAS/STING pathway by NiV and MeV. Our data suggest that cGAS/STING activation is critical in controlling paramyxovirus infection, and possibly represent attractive targets to develop countermeasures against severe disease induced by these pathogens.

## INTRODUCTION

The innate immunity represents the first line of host defense against invading pathogens (Akira et al., 2006). Exogenous motifs associated with viral infections involved in stimulating innate responses include pathogen-derived nucleic acids, DNA or RNA (Akira et al., 2006; Mogensen, 2009). Its ensuing detection activates pattern recognition receptor (PRR)-associated adaptor molecules that are responsible for subsequent expression of type I and III interferons (IFNs) (Park and Iwasaki, 2020) and the induction of IFN-related genes, which are important for the control of virus infection (Borden et al., 2007; Der et al., 1998; Sen and Peters, 2007). Four major axes, defined by their nodal adaptor, are able to induce strong innate immune responses upon sensing of pathogen-related nucleic acids (Baccala et al., 2009). Three of them, Toll-like receptor (TLR)-associated adaptor molecules: myeloid differentiation primary response 88 (MyD88) (Wesche et al., 1997), Toll/interleukin-1 receptor/resistance [TIR] domain-containing adaptor-inducing IFN-β (TRIF) (Yamamoto et al., 2003) and RIG-I-like receptor (RLR)-associated mitochondrial antiviral signaling protein (MAVS) (Seth et al., 2005) are dedicated to sense DNA and/or RNA. The cyclic guanosine monophosphate-adenosine monophosphate (cGAMP) synthase (cGAS)/stimulator of interferon genes (STING also known as ERIS, MITA or TMEM173) pathway is the leading sensor for the detection of cytosolic DNA (Ishikawa and Barber, 2008; Sun et al., 2013). The cGAS/STING axis seems to be involved in the sensing of various different RNA viruses, which may express viral proteins that counteract cGAS/STING activity at different levels. Thus, the cGAS/STING pathway may also contribute to the control of RNA viruses (Ni et al., 2018). Recently, activation of the cGAS/STING pathway has been observed following infection by the positive strand RNA virus SARS-CoV-2 (Neufeldt et al., 2020).

In recent years, members of the *Paramyxoviridae* family have caused numerous emerging zoonoses and/or epidemics (Thibault et al., 2017). This viral family contains both old and new human and zoonotic viral pathogens such as measles virus (MeV) and Nipah virus (NiV). While MeV has been almost eradicated from most developed countries through vaccination campaigns, the number of cases and deaths significantly increased within the last decade killing more than 100.000 people every year (Ferren et al., 2019). Moreover, as NiV is the most virulent paramyxovirus with mortality rates between 40-100% during epidemics and remains without any licensed treatment or vaccine (Pelissier et al., 2019; Soman Pillai et al., 2020), the World Health Organization included it in its blueprint for priority pathogens (Mehand et al., 2018).

Our previous work evaluating innate sensors involved in the protection of mice following NiV infection suggests control of the virus through the activation of MyD88 and MAVS pathways, while TRIF seems dispensable (Iampietro et al., 2020). Although, both MyD88 and MAVS are important to produce high levels of type-I IFNs (IFN-I), double KO mice still exhibit some resistance against NiV infection. This is in contrast to the interferon-α/β receptor (IFNAR) KO mice that lack any IFN-I-related responses and are unable to control NiV infection (Dhondt et al., 2013; Iampietro et al., 2020). We thus hypothesized that the cGAS/STING axis of the innate immunity could also contribute to the control of NiV infection.

Once activated in the cytoplasm of infected cells, cGAS uses adenosine triphosphate (ATP) and guanosine triphosphate (GTP) as substrates for cyclisation into cGAMP. cGAMP triggers STING and further results in IFN-I expression (Ishikawa and Barber, 2008; Sun et al., 2013). Successive conformational changes ensure the activation of cGAS and STING in cascade. Briefly, cGAS in “resting” state is a monomer containing a conserved zinc-ion-binding domain as DNA binding module (Civril et al., 2013; Kranzusch et al., 2014). Upon DNA binding in the cytoplasm, cGAS dimerizes and further oligomerizes to induce optimal cGAMP production (Gao et al., 2013; Zhang et al., 2014). Thereafter, cGAMP binds to STING in its ligand-binding domain and induces inward rotations leading to dimerization and later oligomerization of STING (Cong et al., 2019; Ergun et al., 2019; Shang et al., 2012). In addition, upon binding of cGAMP, STING exits the endoplasmic reticulum to the Golgi apparatus. There, it transduces a downstream signaling pathway by recruiting TANK-binding kinase 1 (TBK1) and IRF-3 (but not IRF-7) transactivator and activating the NF-kB pathway that results in the activation of type I IFN and cytokine genes (Dobbs et al., 2015; Gui et al., 2019; Liu et al., 2015). The recruitment of IRF-3 relies on the phosphorylation of its Ser366 (Ser365 in murine) STING by TBK1 (Tsuchiya et al., 2016). In addition, the E3 ubiquitin ligases TRIM32 and TRIM56 promotes the non-degradative K-63 linked ubiquitination of Lys224 of STING to trigger a cytokine response (Ni et al., 2017; Tanaka and Chen, 2012). As a third type of post-translational modifications (PTMs) STING palmitoylation governs its trafficking to the Golgi (Mukai et al., 2016).

We report here that both cGAS and STING are required for mounting an efficient innate immune response upon NiV and MeV infection. In infected cells, STING is phosphorylated on S366 (S365 in mouse) and is K63 linked ubiquitinated, confirming the presence of its activated form at later time points of RNA virus infection, similarly to what has been observed by others after infection with DNA virus (Chiang and Gack, 2017).

## RESULTS

### STING plays a role in the control of NiV infection in mice

Our previous study revealed a complementary role of MyD88 and MAVS in the partial containment of NiV, suggesting that the complete resistance of mice against NiV involves an additional activation pathway of the IFN-I response (Iampietro et al., 2020). The cGAS/STING axis has recently emerged as critical in the crosstalk between innate sensing of cytosolic DNA and RNA viruses (Ni et al., 2018). We therefore analyzed the susceptibility of mice bearing gene deletions in either single TLR and IL-1R (MyD88 KO) pathway, or in combination with TRIF (MyD88/TRIF KO), MAVS (MyD88/TRIF/MAVS KO) and STING (MyD88/TRIF/MAVS/STING KO) signaling platforms (**Figure 1** **and S1**). The animals were infected intraperitoneally and monitored during 24 days for clinical signs and/or death. While infected wild-type (WT) mice, MyD88 KO and MyD88/TRIF KO mice did not manifest any clinical signs of disease, triple MyD88/TRIF/MAVS KO and quadruple MyD88/TRIF/MAVS/STING KO mice exhibited symptoms of neurological disorders similar to those observed in IFNAR KO mice (**Figure 1A**). Moreover, while 60% triple KO mice survived NiV challenge, all mice bearing the quadruple deficiency succumbed by day 11 post-infection, indicating a crucial and non-redundant role for STING in the protection of mice (**Figure 1A**). NiV nucleoprotein (NiV-N) protein levels in the brain of autopsied animals deficient for MyD88/TRIF/MAVS and MyD88/TRIF/MAVS/STING were comparable to those observed in the brain of IFNAR KO mice at time of death (**Figure 1B**). Furthermore, while analysis of NiV load in murine brain determined equivalent NiV-N RNA levels within these three deficient murine models, evaluation in the spleen showed higher viral loads in mice bearing the quadruple deficiency compared to the triple KO mice (**Figure 1C** **and S1A**). In parallel, the lower production of IFNβ (**Figure 1D** **and S1B**) and IFNα (**Figure 1E** **and S1C**) in the brains and spleens of MyD88/TRIF/MAVS KO and MyD88/TRIF/MAVS/STING KO mice compared to WT was associated with their inability to clear the virus. Overall, these results confirm our previous observations on the importance of the RLR signaling platform involving MAVS (Iampietro et al., 2020) and suggest a novel synergistic and non-redundant role of the STING pathway during NiV infection.

**Figure 1.**
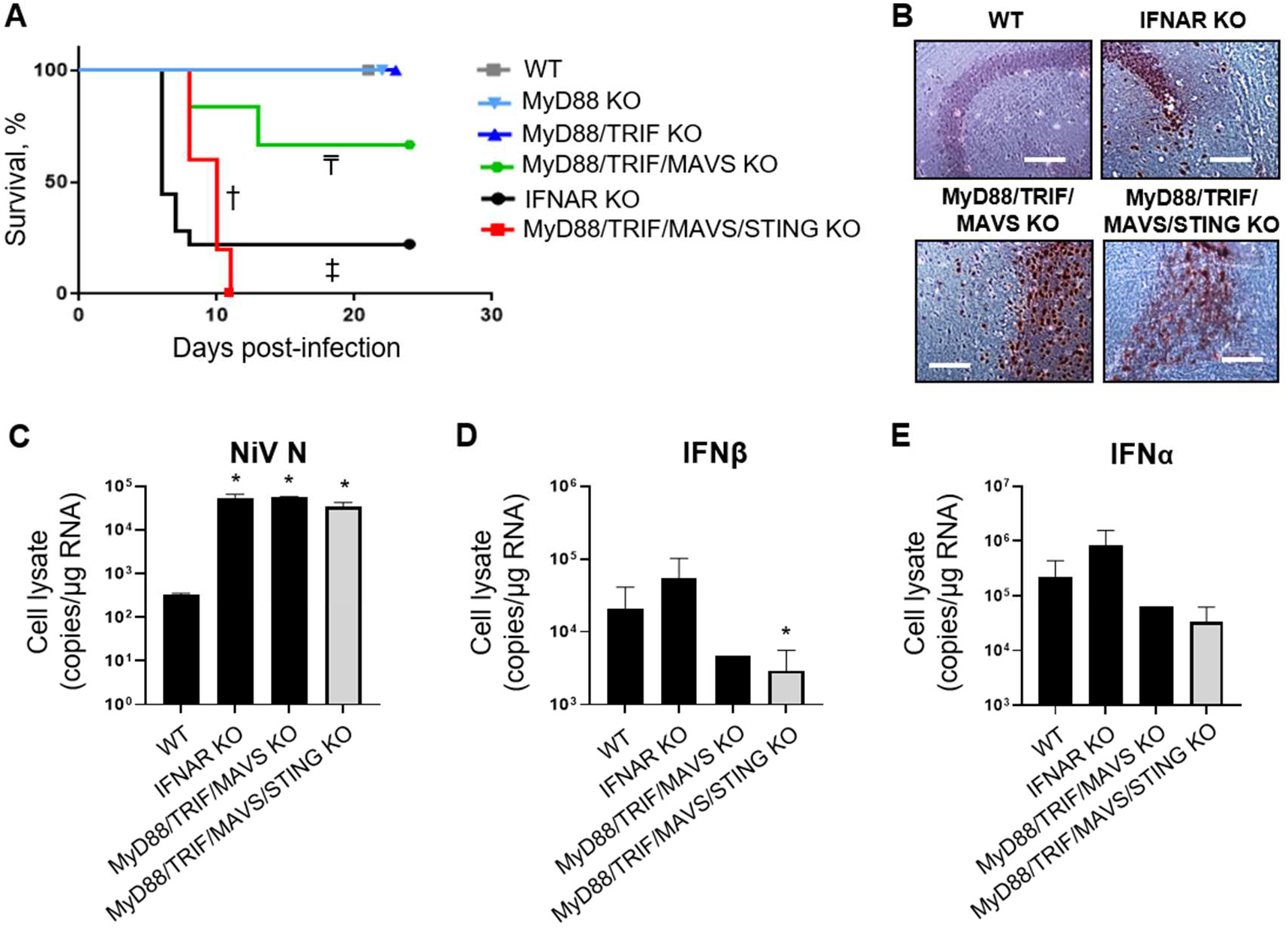
STING plays a role the control of NiV infection in mice. Wild-type (WT) mice and mice deficient in indicated pathogen recognition signaling pathways were infected intraperitoneally with 10^6^ PFU of NiV (5 or 6 animals per group). (A) Survival of mice infected by NiV was followed up for 24 days. ^†^P < 0.05 (MyD88/TRIF/MAVS/STING KO vs WT), ^‡^P< 0.05 (IFNAR KO vs WT) and ^₸^P < 0.01 (MyD88/TRIF/MAVS KO vs MyD88/TRIF/MAVS/STING KO) (Gehan-Breslow-Wilcoxon test). (B) Immunohistochemistry of murine brains following NiV infection. Brains of WT mouse, IFNAR KO mouse, MyD88/TRIF/MAVS KO and MyD88/TRIF/MAVS/STING KO were collected on days 2, 6, 13 and 11 respectively. Scale bars represent 100 μm. (C-E) Expression of NiV nucleoprotein (NiV-N) in the brain of NiV-infected mice, harvested on the day of death or euthanized at the end of protocol for different genotypes, was determined by RT-qPCR. Results represent mean and standard errors for each group. Analysis of IFNβ and IFNα expression by RT-qPCR in organs harvested 2–13 days after infection. All samples were analyzed using One-way analysis of variance, followed by the Tukey multiple comparisons test, \**P*< 0.05 compared to WT condition.

### STING controls NiV replication in primary murine embryonic fibroblasts (pMEFs)

To further evaluate the role of STING during NiV infection in mice, we analyzed the impact of the gene deletion of various nodal adaptors on the *in vitro* NiV replication in pMEFs. The pMEFs were generated from mice bearing the corresponding deleted genes as described previously (Brune et al., 2001). They were infected with rNiV-eGFP to allow imaging of the viral infection by fluorescent microscopy. In agreement with the above *in vivo* observations, MyD88 KO and MyD88/TRIF KO pMEFs were able to control NiV replication as well as WT pMEFs, with only few observed infected cells. In contrast, NiV rapidly spreads within the culture of MyD88/TRIF/MAVS KO and MyD88/TRIF/MAVS/STING KO pMEFs although not as extensively as in IFNAR KO cells (**Figure 2A**). Moreover, single STING KO pMEFs were also unable to control the viral spread, highlighting a major role of STING in the mouse innate defense against NiV infection (**Figure 2A**). The poor permissiveness of WT, MyD88 KO and MyD88/TRIF KO cells was confirmed by a low amount of viral RNA in these cells (**Figure 2B**) and limited release of viral RNA in the supernatant (**Figure 2C**). These three cell types exhibited comparable amounts of IFNβ mRNA (**Figure 2D**) while this mRNA was nearly undetectable in non-infected WT cells. However, contrary to WT pMEFs which exhibit a significant increase in IFNα, the infection of MyD88 KO and MyD88/TRIF KO cells did not result in a significant accumulation of IFNα mRNA (**Figure 2E**). The infection of IFNAR KO cells was associated with elevated cytoplasmic and released viral RNA as well as elevated levels of both IFNβ and IFNα mRNA as expected since all four induction axes of innate immunity are functional but unable to activate an efficient antiviral program (**Figure 2B-E**). In contrast, no accumulation of either IFNβ or IFNα occurred concurrently to the high levels of NiV-N RNA (**Figure 2B-C**) observed in infected MyD88/TRIF/MAVS KO and MyD88/TRIF/MAVS/STING KO (**Figure 2D-E**). In STING KO pMEFs, despite lower levels than IFNAR KO cells, high amounts of NiV-N RNA accumulated in the cytoplasm compared to WT cells (**Figure 2B**). This intermediate phenotype of STING KO cells was associated with no significant IFNβ mRNA induction, but high levels of IFNα mRNA (**Figure 2D-E**). Altogether, these results indicate that in murine pMEFs (i) the control of NiV infection is mediated by IFN-I-mediated activation of an antiviral program, (ii) the IFNβ and IFNα response plays a major and minor role, respectively, in this process, (iii) STING is necessary for the induction of IFNβ, but not of IFNα, and (iv) MAVS, in combination with MyD88 or not, is important for the activation of IFN-I genes confirming our previous observations (Iampietro et al., 2020).

**Figure 2.**
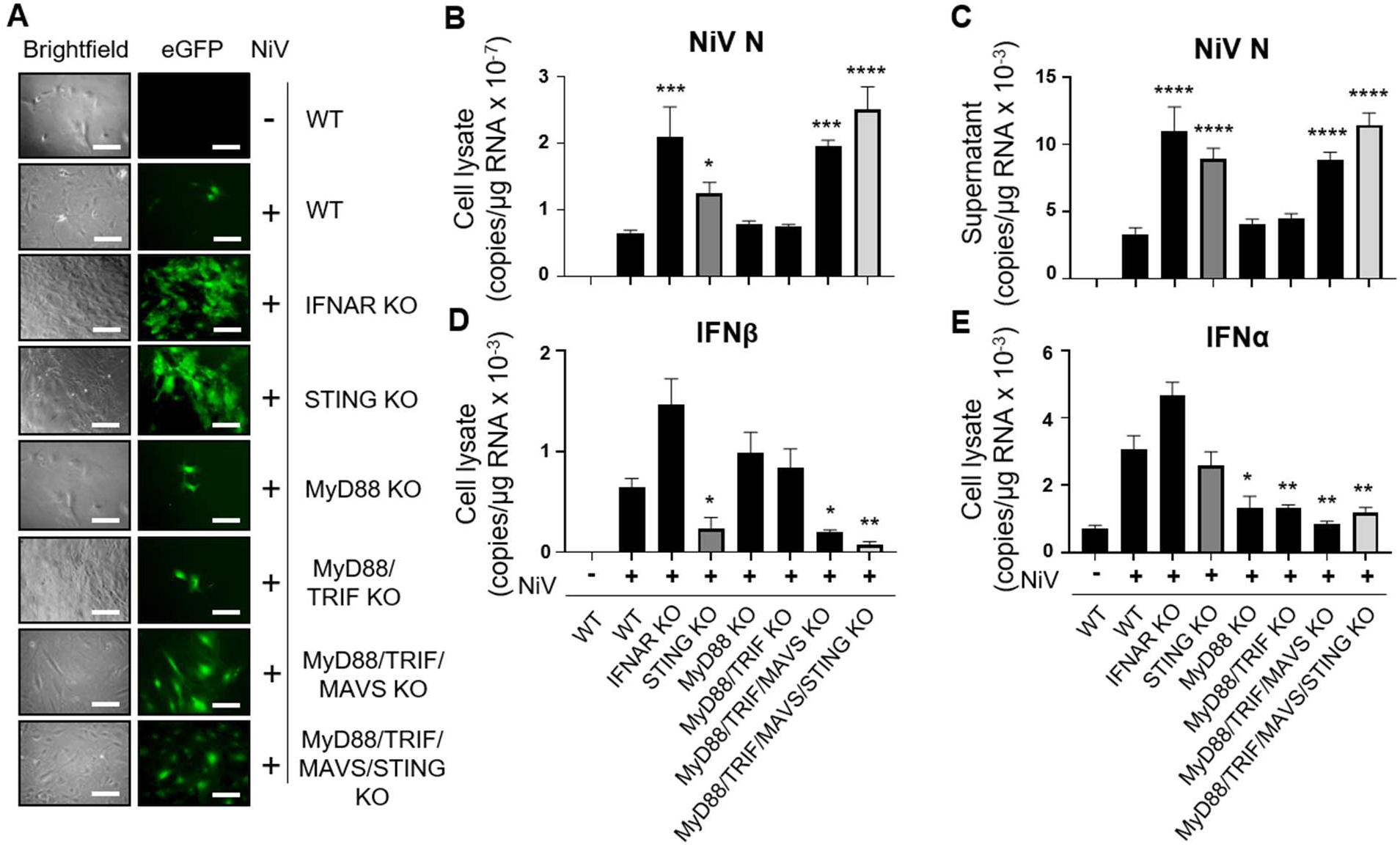
STING controls NiV replication in primary murine embryonic fibroblasts (pMEFs). pMEFs obtained from mice deficient in the indicated signaling pathways were infected with rNiV–eGFP (MOI of 0.3) and cultured for 24 h. (A) Cells were analyzed for eGFP expression by fluorescence microscopy. Scale bars represent 100 μm. (B–E) Cells and supernatants were harvested and analyzed by RT-qPCR for NiV-N (B and C), IFNβ (D), and IFNα (E) expression. Results are presented as means ± standard errors. The statistical significance of differences between infected wild-type (WT) cells and knockout (KO) cells was analyzed using 1-way analysis of variance, followed by the Tukey multiple comparisons test. \**P* < 0.05; \*\**P* < 0.01; \*\*\**P* < 0.001; \*\*\*\**P* < 0.0001 compared to NiV-infected WT condition.

### cGAS/STING pathway has a critical role in the control of paramyxovirus infection in human THP-1 cells

The role of STING and its upstream activator cGAS in the innate response to NiV was further explored in the human monocytic THP-1 cell line that was either WT or deficient in cGAS (cGAS KO) or STING (STING KO). WT THP-1 cells have the advantage to exhibit a limited permissiveness to infection with both NiV and the WT derived recombinant MeV strain rMeV-EdmH-eGFP another member of the *Paramyxoviridae* family belonging to the closely related *morbillivirus* genus. This virus expresses the Edmonston vaccine-derived H glycoprotein allowing the use of ubiquitously expressed CD46 as cellular receptor (Naniche et al., 1993) and hence enters into THP-1 cells that do not or minimally express the known physiological MeV receptors CD150 and Nectin 4 (Crimeen-Irwin et al., 2003; Noyce et al., 2011; Tatsuo et al., 2000). THP-1 cells depleted of cGAS or STING exhibited enhanced permissiveness to NiV and MeV infections when compared to their WT counterparts (**Figure 3A, 3B**, **S2A and S2B**). This was analyzed by fluorescence imaging of NiV propagation throughout the cell culture (**Figure 3A**), by the proportions of THP-1 infected cells as determined by flow cytometry (**Figure S2A**), and by the quantification of viral N RNA by RT-qPCR both in cell extracts (**Figure 3C**) and released in the cell supernatants (**Figure 3D**). Correlatively, and in agreement with observations made in pMEFs infected with NiV, the higher permissiveness of cGAS KO and STING KO THP-1 cells was associated with the abolition of IFNβ and/or IFNα mRNA accumulation (**Figure 3E-F**). Comparable results were obtained after infection by MeV although a reduced but significant accumulation of both IFNβ and IFNα mRNA was still observed in cGAS KO and STING KO cells (**Figure 3G-J).**

**Figure 3.**
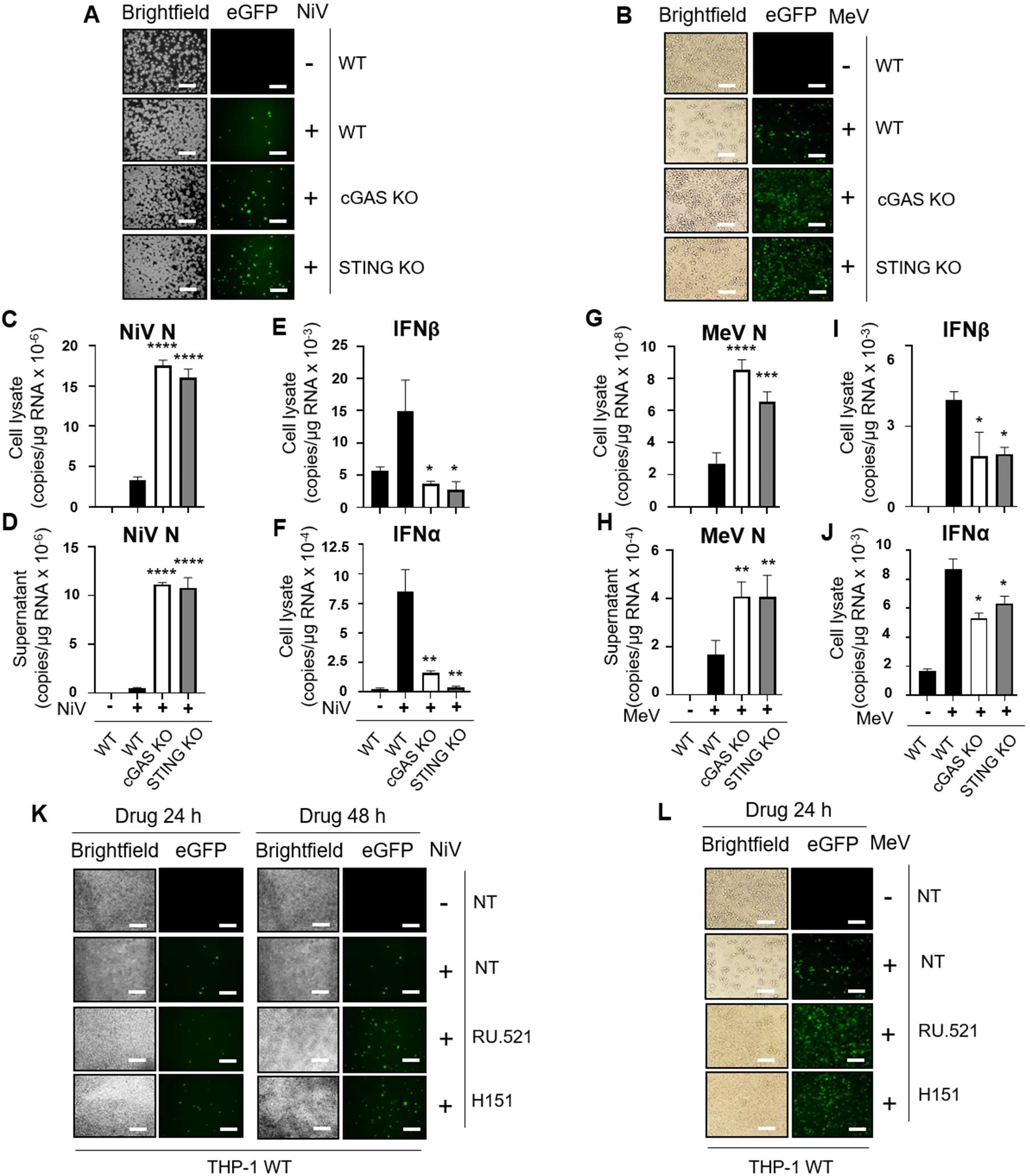
cGAS/STING pathway has a critical role in the control of paramyxovirus infection in human THP-1 cells. THP-1 cells deficient in the indicated signaling pathways were infected with NiV–eGFP and MeV-eGFP (MOI of 0.1) for 48 h and 24 h respectively. (A-B) Cells were analyzed for eGFP expression by fluorescence microscopy. (C-J) Cell lysates and/or supernatants were harvested and analyzed by RT-qPCR for the expression of NiV-N (C and D), MeV-N (G and H), IFNβ (E and I), and IFNα (F and J). Results are presented as means ± standard errors. THP-1 WT cells treated or not (NT) with the specific inhibitors for cGAS (RU.521) or STING (H151) were infected with NiV–eGFP and MeV-eGFP (MOI of 0.1) for 24 h and 48 h. (K and L) Cells were analyzed for eGFP expression by fluorescence microscopy. Scale bars represent 100 μm. The statistical significance of differences between infected WT and KO cells was analyzed using 1-way analysis of variance, followed by the Tukey multiple comparisons test. \**P*< 0.05; \*\**P* < 0.01; \*\*\**P*< 0.001; \*\*\*\**P* < 0.0001 compared to infected WT condition.

As an alternative and complementary approach, we analyzed the paramyxoviral propagation within WT THP-1 cells treated with selective inhibitors of cGAS and STING, namely RU-521 and H151 drugs, respectively. RU.521 is a competitive inhibitor of ATP and GTP substrates for binding to the cGAS catalytic pocket that prevents their cyclisation into cGAMP (Vincent et al., 2017; Xie et al., 2019). H151 covalently binds to STING Cys91, inhibits the palmitoylation of STING and consequently the activation of IFN-I production (Haag et al., 2018). Addition of RU-521 or H151 one hour prior to the infection resulted in an improved propagation of NiV and MeV as evidenced by fluorescence microscopy (**Figure 3K** **and** **3L**) and flow cytometry (**Figure S2C, S2D, S2E**).

Thus, the cGAS/STING axis also appears to play a critical role in human cells to control paramyxovirus infections by allowing the expression of IFNβ and to a lower extent that of IFNα.

### Paramyxovirus infection activates the cGAS/STING pathway in both murine and human cells

The increased viral infection observed upon abolition of either cGAS or STING suggested that paramyxoviruses could activate STING. This was investigated by analyzing specific phosphorylation and/or ubiquitination of activated STING in pMEFs, THP-1 and human pulmonary microvascular endothelial cells (HPMEC) as representative of primary targets of NiV infection in humans (**Figure 4** **and S3**). A time course follow up of the activation of cGAS/STING axis induced by viral infection was performed at 6, 24 and 48 hours post-infection (h.p.i.) by western blot using antibodies against STING-S366^P^ (Chiang and Gack, 2017). While the expression level of STING remained unchanged throughout the 48h observation, moderate amounts of STING-S366^P^ became detectable at 24 h.p.i. and further increased at 48 h.p.i. after NiV or MeV infection of HPMECs (**Figure 4A** **and** **4B**) and THP-1 cells (**Figure 4C** **and S3A**). Importantly, STING-S366^P^ was not detected in NiV- or MeV-infected cGAS KO THP-1 cells at 24 and 48 h.p.i., and as expected in STING KO THP-1 cells (**Figure 4C** **and S3A**). Accordingly, STING-S366^P^ was also not detected in THP-1 cells infected and treated with either RU-521 or H151 (**Figure S3A**).

**Figure 4.**
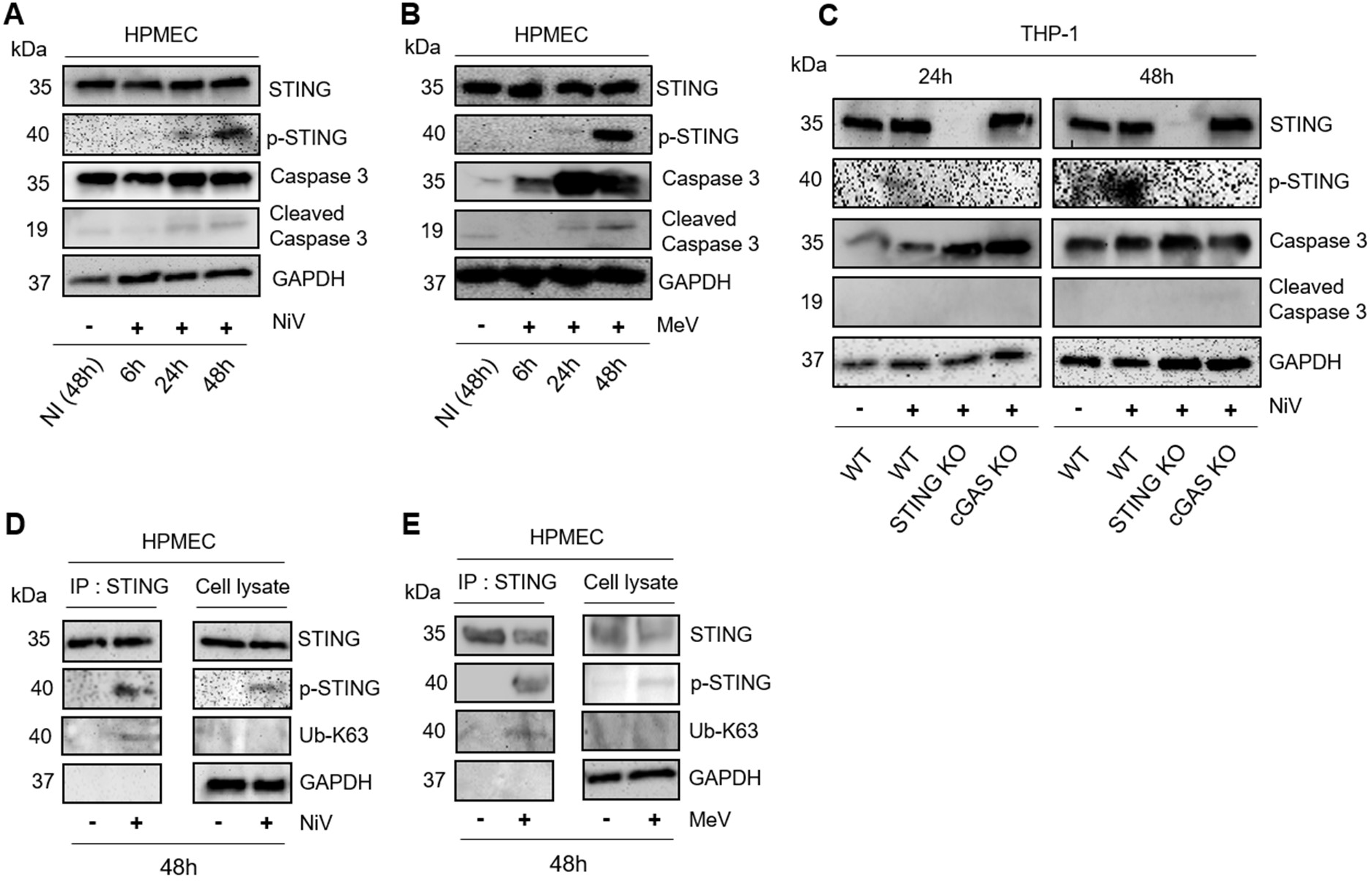
Paramyxovirus infection activates the cGAS/STING pathway in human cells. Human pulmonary microvascular endothelial (HPMEC) cells and THP-1 cells deficient in the indicated signaling platforms were infected with either NiV or MeV (MOI of 1) for 6, 24 or 48 h. (A-C) Cells were analyzed for phospho-STING (p-STING), STING, Caspase 3, cleaved Caspase 3 and GAPDH expression by western blot analysis. (D–E) HPMEC cells were infected with NiV (D) and MeV (E) (MOI of 1) for 48 h before cell lysis. Endogenous STING was immunoprecipitated using anti-STING antibodies followed by western blot analyses using anti-STING, anti-Ub-K63, anti p-STING and anti-GAPDH antibodies. In parallel, endogenous expression levels of STING, p-STING and Ub-K63 together as GAPDH as loading control in cell lysates were analyzed by western blot. Shown data are representative of three independent experiments showing similar results.

Accordingly, while uninfected WT pMEFs, NiV-infected STING KO and MyD88/TRIF/MAVS/STING KO cells did not elicit any mouse STING-S365^P^ band (Tsuchiya et al., 2016), NiV-infected WT and MyD88/TRIF/MAVS KO pMEFs displayed activation of STING protein (**Figure S3B**). Importantly, the detection of the STING-S365^P^ band in MyD88/TRIF/MAVS KO pMEFs indicates that NiV infection activates the STING signaling pathway independently from the TLR-MyD88/TRIF and RLR/MAVS pathways of innate immunity. Since K63-linked ubiquitination (Ub-K63) is another hallmark of STING activation, mainly associated to the activation of NF-κB (Chiang and Gack, 2017), STING ubiquitination was also evaluated in HPMECs infected for 48 h with NiV or MeV. STING immunoprecitated using anti-STING antibodies was found to be labelled by both anti-STING-S366^P^ and anti-Ub-K63 antibodies (**Figure 4D** **and** **4E**). As an apoptotic environment could activate STING in bystander cells (Ahn et al., 2012), a potential effect of viral-induced cell death in infected cultures was evaluated by analyzing caspase 3 activity. The levels of cleaved caspase 3 were minimally increased between uninfected and NiV- or MeV-infected cells, indicating that cell death had likely a minor impact on STING activation until 48 h post-infection (**Figure 4A, 4B**, **S3A, S3B, S3C and S3D**).

We conclude that negative strand RNA paramyxoviruses activate the cGAS/STING axis to trigger the innate immune responses.

## DISCUSSION

Although its role in response to DNA virus infection has been well deciphered (Eaglesham and Kranzusch, 2020), the role of cGAS/STING against RNA viruses such as paramyxoviruses is poorly understood. While cGAS was reported to interact with dsRNA, no production of cGAMP was detected, thus cGAS could not canonically activate STING directly (Civril et al., 2013). However, previous studies have shown interactions between RNA viruses and cGAS/STING in (i) their control by innate immunity or (ii) their capacity to disrupt the cGAS/STING pathway, reflecting a likely role of STING in the host response(Aguirre et al., 2017; Franz et al., 2018; Ishikawa et al., 2009; Schoggins et al., 2014).

As paramyxoviruses replicate within the cytoplasm using their own RNA-dependent RNA polymerase, i.e. their replicative cycle does not rely on a DNA intermediate step (Gerlier and Lyles, 2011). Because cGAS is exclusively activated upon binding to dsDNA (Kranzusch et al., 2013; Sun et al., 2013) or DNA/RNA hybrid (Mankan et al., 2014), we have to speculate about the identity of the cGAS agonist during infection by paramyxoviruses. The possibility of mitochondrial release of DNA at some stage of the viral infection as evidenced after infection with Dengue virus, a positive stranded RNA virus (Aguirre et al., 2017; Sun et al., 2017), will have to be explored.

Collectively, the obtained data fit with a model where the infection by NiV or MeV activates, the cGAS/STING pathway. This activation results in the induction of the IFNβ and possibly the IFNα gene (see below) that mediate the activation of an efficient antiviral program in *in vitro* and *in vivo* mouse models and/or in human cells closely mimicking what happens after the infection by mouse cytomegalovirus (MCMV), a DNA virus (Tegtmeyer et al., 2019). Notably, the comparison of the phenotype of triple and quadruple KO pMEFs indicate that the cGAS/STING pathway strongly reinforces (or even conditions) the activation of type I IFN responses via the TLR8/MyD88 and/or the RLR/MAVS pathway by paramyxoviruses, including MeV (Ikegame et al., 2010; Runge et al., 2014; Seth et al., 2005).

STING can bind to RIG-I and MAVS (Sun et al., 2012) and is thought to potentiate the RLR/MAVS signaling leading to the activation of type I IFN (Ishikawa et al., 2009; Zevini et al., 2017; Zhong et al., 2008). The present data fits with this model. The defect of the cGAS/STING pathway strongly facilitates the replication of NiV and MeV as expected from the observed loss in activating the IFN-I response. We cannot exclude that this facilitation results also from the concomitant disappearance of a STING-dependent inhibition of the translation of viral protein as recently reported (Franz et al., 2018). However, they reported that the absence of STING modestly affects the IFN response to the infection by negative stranded RNA viruses compared to the present work.

Upon infection of MeV and/or NiV in mouse and/or human cells, the absence of the cGAS/STING pathway affect the IFNβ response and to a lower and more variable extend the IFNα response. Due to cell-dependent ability to produce IFNα, different outcome may reflect higher expression of IRF-7 in pMEFs (Sharma et al., 2019) compared to THP-1 cells (Green et al., 2020). Indeed, while the activation of the IFNβ gene optimally relies on IRF3 homodimers or IRF3/IRF7 heterodimers and NF-κB (Honda et al., 2005; Wathelet et al., 1998) that of the IFNα genes relies mostly on IRF7 homodimers (Yeow et al., 2000). Interestingly, STING mostly targets IRF-3 (Zhong et al., 2008) and NF-κB (Stempel et al., 2019) and consequently STING preferentially activates IFNβ.

Our study demonstrates that STING is activated against RNA viruses as also recently reported with SARS-CoV-2 (Neufeldt et al., 2020) and highlights that STING is modified and activated through S366 phosphorylation and/or K-63 linked ubiquitination during NiV and MeV infection. Additional studies will uncover the occurrence of others PMTs modification, STING subcellular location and the source of the cognate DNA that activate cGAS during RNA virus infection.

In conclusion, cGAS/STING activation occurs during paramyxovirus infections, both *in vitro* and *in vivo*. This highlights an undefined aspect of the immune regulation against negative strand viruses and reveals cGAS/STING as potential targets in the development of novel antiviral strategies.

## METHODS

### Mice

Several lines of transgenic mice, all in C57BL/6 genetic background, were used: wild type (WT) C57BL/6J mice and following knockout (KO) models: mice deleted for IFN-I receptor (IFNAR-KO) (Muller et al., 1994), TLR adaptor protein MyD88 (MyD88-KO) (Adachi et al., 1998), and mice crossed to bear several deletions, including MyD88/TRIF-KO (Waibler et al., 2007), MyD88/TRIF/MAVS-KO (Spanier et al., 2014) and MyD88/TRIF/MAVS/STING KO (Tegtmeyer et al., 2019).

### Infection of mice

Groups of 5–10 mice from each line, 4–6 weeks of age, were anesthetized with isoflurane and infected intraperitoneally with 10^6^ PFUs of Nipah virus Malaysia strain, contained in a 200 μL volume. Animals were monitored for 24 days after infection and manipulated in accordance to good experimental practice and approved by the regional ethics committee CECCAPP (Comité d’Evaluation Commun au Centre Léon Bérard, à l’Animalerie de transit de l’ENS, au PBES et au laboratoire P4) and authorized by the French Ministry of Higher Education and Research (no. 00962.01). Animal experiments were conducted by the animal facility team in the INSERM Jean Mérieux BSL-4 laboratory in Lyon, France.

### Cell Lines

Primary murine embryonic fibroblasts (pMEFs) were isolated from murine embryos obtained from pregnant mice 13 days after conception, as described elsewhere (Brune et al., 2001) and cultured in Dulbecco’s modified Eagle’s medium (DMEM) GlutaMAX supplemented with 10% heat-inactivated fetal bovine serum (FBS), 0.2% 2-mercaptoethanol, 1% HEPES, 1% nonessential amino acids, 1% sodium pyruvate, and 2% penicillin-streptomycin mix. For infections, pMEFs were plated in 12-well plates at 2.10^5^ cells per well and cultured with rNiV-eGFP at a multiplicity of infection (MOI) of 0.3 plaque-forming units (PFUs)/cell for 1 h at 37°C. Virus-containing medium was then removed, and cells were washed once with 1× phosphate-buffered saline. Finally, fresh DMEM was added to cells that were incubated for 24 h at 37°C. Human monocytic THP-1 cell lines were obtained as described previously and cultured in RPMI 1640 GlutaMAX supplemented with 10% heat-inactivated FBS, 1% HEPES and 2% of penicillin-streptomycin mix. Human pulmonary microvascular endothelial cells (HPMEC) (Krump-Konvalinkova et al., 2001) were cultured in Endothelial Cell Growth Medium, in flasks coated with 0,1% bovine gelatine in PBS. For infection, HPMECs were plated in 12-well plates at 2.10^5^ cells per well and cultured with rNiV-eGFP or rMeV-edmH-eGFP at a MOI of 1 PFUs/cell for 24 or 48 hours at 37°C. All cell types were incubated at 37°C with 5% CO2 and were tested negative for Mycoplasma spp.

### Drugs

H151, a specific inhibitor for STING and RU.521, a specific inhibitor for cGAS were added 1 h previous infection of THP-1 cells at 10 μM and 10 μg/ml, respectively, selected according to the previously published results (Chang et al., 2020; Hayden et al., 2020). Then, cells were infected with rNiV-eGFP or rMeV-edmH-eGFP and incubated for 24 or 48 h at 37°C with 5% CO2.

### Viruses

NiV Malaysia (isolate UMMC1; GeneBak AY029767, recombinant NiV (rNiv)–enhanced green fluorescent protein (eGFP) (Yoneda et al., 2006) and recombinant MV IC323 vaccine strain, expressing Edmonston H and eGFP (Hashimoto et al., 2002), kindly provided by Dr Y. Yanagi (Kyushu University, Japan) and were prepared by infecting Vero-E6 cells, in the INSERM Jean Mérieux biosafety level 4 (BSL-4) and BSL-2 laboratories at CIRI in Lyon, France respectively.

### Immunohistochemistry

Brains from mice were embedded in paraffin wax and sectioned at 7 μm. Slides were deparaffinated and rehydrated in three Xylene baths for 5 min each, followed by two 100% alcohol baths for 5 min, and then succeeded with multiple baths using decreasing level of alcohol for 3 min each. After deparaffination, slides were put in a sodium citrate solution in a boiling water bath for 20 min for heat-induced epitope retrieval and washed 3 times in PBS for 3 min afterwards. Activity of endogenous peroxydase was blocked using a H2O2 0.3% solution. Blocking of non-specific epitopes is done using PBS-2.5% decomplemented Normal Horse Serum + 0.15% Triton X-100 for 30 min. Then, primary rabbit anti-NiV N antibody was used at 1/10000 dilution and incubated overnight at 4°C in the blocking buffer. For secondary antibody and revealing steps, ImmPress system (anti-rabbit ig/peroxydase) was used. Counterstaining was performed using Harris solution and photographs were taken with a microscope Zeiss Axiovert 100M.

### Co-immunoprecipitations

HPMEC cells (5×10^5^) were seeded in 6-well plates. 16 h after seeding, cells were infected with the appropriate dilution of rNiV-eGFP or rMeV-edmH-eGFP at a MOI of 1 in RPMI described above. Forty-eight hours post-infection, cells were lysed in RIPA buffer, supplemented with a cocktail of protease-phosphatase inhibitors for 30 min on ice, and centrifuged for 10 min at 4°C at 15,000 g. Supernatants were incubated with a rabbit anti-STING antibody for 2 h at 4°C. Then, protein A/G agarose beads were added to the mix overnight at 4°C. Beads were then washed three times in washing buffer (RIPA buffer, supplemented with a cocktail of protease-phosphatase inhibitor), and proteins were eluted in 100 μl of elution buffer (Reducing agent 10X, Laemmli 4X, RIPA buffer, supplemented with a cocktail of protease-phosphatase inhibitors) for 15 min at 96°C. Then the eluate and a sample of input of the cell extract were run on polyacrylamide gel electrophoresis (SDS-PAGE) and analyzed by western-blotting.

### Immunoblot Analysis

Heated protein lysates were separated by 4-15% SDS-PAGE and electro transferred for 1 h onto polyvinylidene difluoride (PVDF) membranes at 4°C. PVDF membranes were blocked in Tris-buffered saline containing 0.05% Tween 20 (TBS-T) + 5% milk for 1 h and then incubated overnight with primary antibodies, mouse anti-GAPDH, rabbit anti-STING, rabbit anti-human S366 p-STING, rabbit anti-mouse S365 p-STING, rabbit anti-Caspase 3, rabbit anti-cleaved Caspase 3 and rabbit anti-Ub-K63 antibodies, diluted 1:1000 in TBS-T + 0.2% milk. Membranes were then washed 3 times using TBS-T and incubated on an additional 1 h with horseradish peroxydase conjugated anti-mouse or anti-rabbit IgG antibodies (diluted 1:5000 in TBS-T + 5% milk). Membranes were then washed 5 times in TBS-T, incubated in Super Signal West Dura to stain cell lysates or in Super Signal West Femto reagent to stain bead eluates. Chemiluminescent signals were measured with the VersaDoc and ChemiDoc Imaging System.

### RNA Extraction and RT-qPCR

At indicated time points, cells and supernatants were collected and RNA extracted using appropriate NucleoSpin RNA Kits according to the manufacturer’s instructions and yield and purity of extracted RNA was assessed using the DS-11-FX spectrophotometer. Equal amounts of extracted RNA (500 ng) were reverse transcribed using the iScript Select cDNA Synthesis Kit and amplified by real-time PCR using Platinum SYBR Green qPCR SuperMix-UDG on a StepOnePlus Real-Time PCR System. Data were analyzed using StepOne software and calculations were done using the 2^ΔΔCT^ method. Expression was normalized to that of glyceraldehyde 3-phosphate dehydrogenase (GAPDH) and expressed as copies of mRNA. Specific set of primers were designed and validated for the detection of human hGAPDH, hIFNα and hIFNβ, murine mGAPDH, murine mIFNα and mIFNβ and viral NiV N and MeV N.

### Immunofluorescence

For PMEFs and THP-1 infections, 3×10^5^ and 5×10^5^ cells were seeded in 12-well plates, respectively, before being infected with rNiV-eGFP and rMeV-edmH-eGFP at a MOI of 0.3 or 0.1 and cultured for 24 or 48 h at 37°C with 5% CO2. Then, cells were evaluated for eGFP expression using a Zeiss Axiovert 100M microscope in the BSL-4 or a NIKON Eclipse Ts2R in the BSL-2, and photographs were taken 24 and 48 h after infection and treated using ImageJ software version Java 1.8.0_112.

### Flow cytometry

THP-1 cells were seeded in 12-well plates at 3×10^5^ and 5×10^5^ per well before being infected with rNiV-eGFP and rMeV-edmH-eGFP at a MOI of 0.1 and cultured for 24 or 48 h at 37°C with 5% CO2. Then, cells were washed, reconstituted in PBS 1X and evaluated for eGFP expression using a Gallios flow cytometer in the BSL-4 or a 4L Fortessa flow cytometer. Analyses were performed 24 and 48 h after infection.

### Ethical statement

Animals were handled in strict accordance with good animal practice as defined by the French national charter on the ethics of animal experimentation and all efforts were made to minimize suffering. Animal work was approved by the Regional ethical committee and French Ministry of High Education and Research and experiments were performed in the INSERM Jean Mérieux BSL-4 laboratory in Lyon, France (French Animal regulation committee N° 00962.01).

### Analysis of eGFP quantification in THP-1 cells

The results are presented in the form of histograms which represent the mean eGFP positive cells for each conditions and error bars represent the standard errors (SE) for n=3 experimental replicates. The different conditions were compared to the control (WT+). Statistical significance was assessed by a one-way ANOVA, followed by a Tukey’s multiple comparisons test; *p < 0.05, **p < 0.01, ***p < 0.001 and ****p < 0.0001 (threshold of significance of 5%).

### qPCR analysis

The results are presented in the form of histograms, which represent the mean of copies of mRNA for a gene for each condition and error bars represent the standard errors (SE) for n=3 experimental replicates. The different conditions were compared to the control (WT+). Statistical significance was assessed by a one-way ANOVA, followed by a Tukey’s multiple comparisons test; *p < 0.05, **p < 0.01, ***p < 0.001 and ****p < 0.0001 (threshold of significance of 5%).

### Densitometry

Densitometric analysis of cleaved caspase 3 immunoblots from three independent experiments were performed using the VersaDoc Imaging System (Bio-Rad) and analyzed with ImageJ 1.52p Fiji package software (https://imagej.net/Fiji). GAPDH expression was used for normalization.

## ACKNOWLEDGEMENTS

The work was supported by INSERM, LABEX ECOFECT (ANR-11-LABX-0048) of Lyon University, within the program “Investissements d’Avenir” (ANR-11-IDEX-0007) operated by the French National Research Agency (ANR), by ANR-18-CE11-0014-02, by Aviesan Sino-French agreement on Nipah virus study. We thank the entire animal experimentation team of INSERM “Jean Mérieux” BSL4 laboratory for the realization of the animal experiment and the biosafety team for their assistance for BSL4 activities. We are grateful to all the members of the group Immunobiology of viral infection at CIRI, for the help in the realization of this study. We acknowledge the contribution of the SFR Biosciences (UMS3444/CNRS, US8/INSERM, ENS de Lyon, UCBL) flow cytometry facility.

## AUTHOR CONTRIBUTIONS

MI, UK and BH designed the study. MI, CD, CM, JR, AC, SD, KD, NA and SM performed experiments. MI, CM, JS, DG and BH analyzed the data. MI, JS, DG and BH wrote the article. MI, JS, DG and BH prepared the figures. MF, RP, JS and UK provided some essential tools.

