## Supplemental Data for "Activation of cGAS/STING pathway upon paramyxovirus infection"

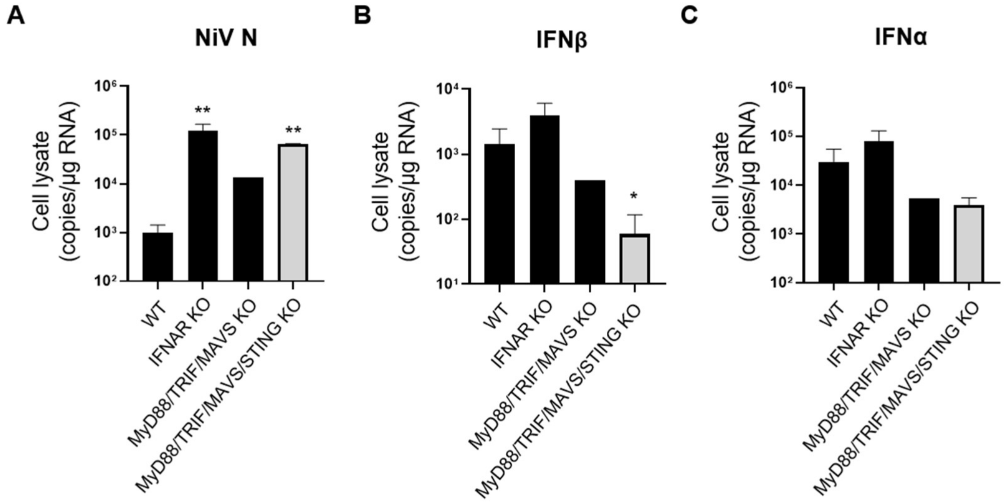


**Figure S1. STING is involved for the control of NiV infection in the murine spleen.** Wild-type (WT) mice and mice deficient in indicated pathogen recognition signaling pathways were infected intraperitoneally with $10^6$ PFU of NiV (5 or 6 animals per group). (A-C) Expression of NiV nucleoprotein (NiV-N) in murine spleen, harvested on the day of death or at the end of protocol for different genotypes, was determined using RT-qPCR. Results represent mean and standard errors for each group. Analysis of IFNβ and IFNα expression by RT-qPCR in organs harvested 2–13 days after infection. All samples were analyzed using One-way analysis of variance, followed by the Tukey multiple comparisons test, **P*< 0.05; ***P*< 0.01 compared to WT condition.

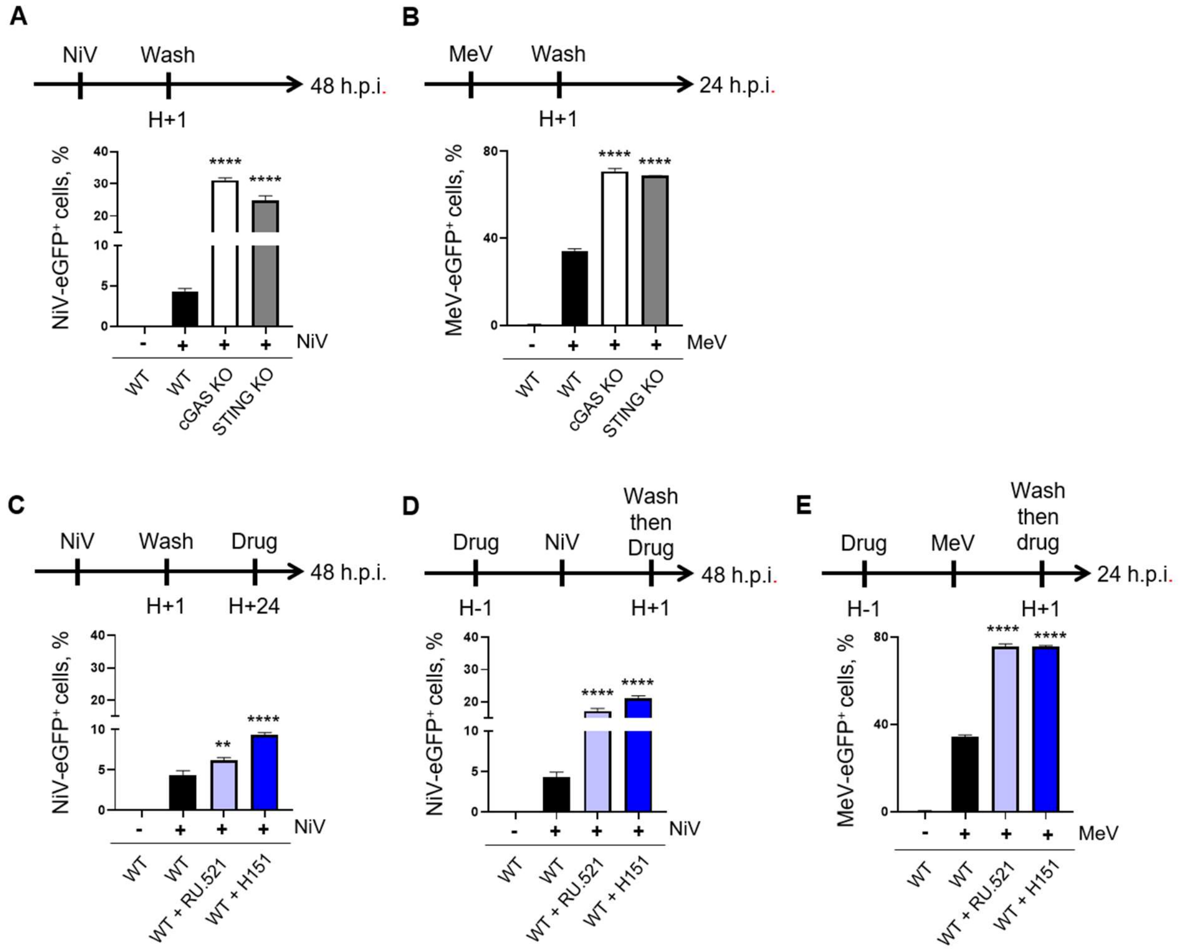


**Figure S2. cGAS and STING signaling control NiV infection in THP-1 cells.** THP-1 cells deficient in the indicated signaling platforms (A, B) or treated with specific inhibitors for cGAS or STING (C-E) were infected with recombinant NiV–eGFP (A, C, D) for 24 and 48 h and MeV-eGFP (B, E) for 24 h (MOI of 0.1) respectively. (A-E) Corresponding experimental procedures are schemed above each result. Cells were analyzed for eGFP expression by fluorescence microscopy and quantified by flow cytometry analysis. All samples were analyzed using One-way analysis of variance, followed by the Tukey multiple comparisons test, $^{**}P < 0.01$; $^{****}P < 0.0001$ compared to infected WT condition.

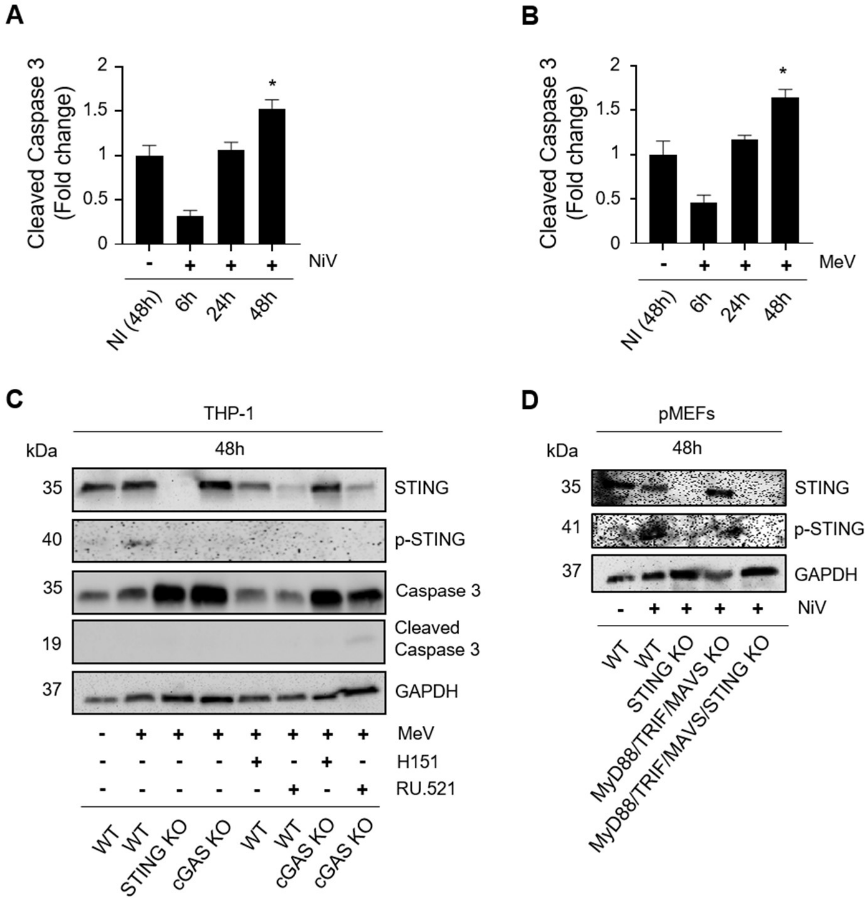


**Figure S3. cGAS/STING pathway is induced by paramyxoviruses in human and murine cells.** (A, B) Densitometric analyses of cleaved caspase 3 in HPMEC cells infected with either NiV (A) or MeV (B) for 6, 24 or 48 h. THP-1 (C) and pMEFs (D) cells deficient in the indicated signaling platforms or treated or not with the specific inhibitors for cGAS (RU-521) or STING (H151) were infected with either MeV (C) or NiV (D) at MOI1 for 48 h. Cells were analyzed for phospho-STING (p-STING), STING, Caspase 3, cleaved Caspase 3 and GAPDH expression by western blot analysis. GAPDH was used as a loading control. All samples were analyzed using One-way analysis of variance, followed by the Tukey multiple comparisons test, **P*< 0.05 compared to infected NI (48h) condition. Data set representative of three independent experiments.
